# Helicobacter pylori-induced adrenomedullin modulates IFN-γ-producing T-cell responses and contribute to gastritis

**DOI:** 10.1101/482125

**Authors:** Hui Kong, Jin-yu Zhang, Fang-yuan Mao, Yong-sheng Teng, Yi-pin Lv, Yu-gang Liu, Weisan Chen, Yu Zhang, Gang Guo, Yuan Zhuang

## Abstract

Adrenomedullin (ADM) is a multifunctional peptide that is expressed by many surface epithelial cells, but its relevance to *H. pylori*-induced gastritis is unknown. Here, we found that gastric ADM expression was elevated in gastric mucosa of *H. pylori*-infected patients and mice. In *H. pylori*-infected human gastric mucosa, ADM expression was positively correlated with the degree of gastritis, accordingly, blockade of ADM resulted in decreased inflammation within the gastric mucosa of *H. pylori*-infected mice. During *H. pylori* infection, ADM production was promoted via PI3K-AKT signaling pathway activation by gastric epithelial cells in a *cagA*-dependent manner, and resulted in increased inflammation within the gastric mucosa. This inflammation was characterized by the increased IFN-γ-producing T cells, whose differentiation was induced via the phosphorylation of AKT and STAT3 by ADM derived from gastric epithelial cells. ADM also induced macrophages to produce IL-12, which promoted the IFN-γ-producing T-cell responses, thereby contributing to the development of *H. pylori*-associated gastritis. Accordingly, blockade of IFN-γ or knockout of IFN-γ decreased inflammation within the gastric mucosa of *H. pylori*-infected mice. This study identifies a novel regulatory network involving *H. pylori*, gastric epithelial cells, ADM, macrophages, T cells, and IFN-γ, which collectively exert a pro-inflammatory effect within the gastric microenvironment.

**Author summary:** *H. pylori* infect almost half the world’s population. Once infected, most of people carry the bacteria lifelong if left untreated, so that persistent *H. pylori* infection can lead to chronic gastritis, peptic ulceration and ultimately gastric cancer. *H. pylori* infection is accompanied with increased inflammation in gastric mucosa, but the mechanisms of chronic gastritis induced by *H. pylori* infection remains poorly understood. We studied a multifunctional peptide known as adrenomedullin (ADM) in gastric epithelial cells, which was known as a key factor of regulating gastrointestinal physiology and pathology. Here, we found that gastric ADM expression was elevated in gastric mucosa of *H. pylori*-infected patients and mice, and was positively correlated with the degree of gastritis. ADM production was promoted via PI3K-AKT signaling pathway activation by gastric epithelial cells in a *cagA*-dependent manner. Blockade of ADM during *H. pylori* infection resulted in decreased gastric inflammation that was characterized by the increased IFN-γ-producing T cells which was induced via the phosphorylation of AKT and STAT3 by ADM derived from gastric epithelial cells. ADM also induced macrophages to produce IL-12, which promoted the IFN-γ-producing T-cell responses. These data demonstrate that *H. pylori*-induced ADM modulates FN-γ-producing T-cell responses and contribute to gastritis.

## Introduction

*Helicobacter pylori (H. pylori)* is a gram-negative bacterium that infects more than half of the world’s population [1]. *H. pylori* is an important factor of chronic gastritis, peptic ulcer and other digestive system diseases, and has been classified as a class I carcinogen by WHO [2]. During the infection, gastric epithelial cells produce a variety of cytokines that are involved in the inflammatory gastric environment after contacting with *H. pylori* [3]. Besides, many immune cells such as neutrophils, lymphocytes and plasma cells are releasing inflammatory factors in the stomach of *H. pylori*-infection [4-6]. Inflammatory reaction to *H. pylori* infection shows special characteristics rarely seen in other organs or biological systems, and the mixed acute and chronic inflammatory reactions contribute to *H. pylori*-associated gastritis and take place simultaneously during *H. pylori* infection.

Adrenomedullin (ADM) is a small active hormone which is expressed throughout the gastrointestinal tract [7]. ADM that consists of 52 amino acids is structurally similar to calcitonin gene-related peptide, dextrin, and pituitary [8]. ADM is abundant in the gastrointestinal tract, especially in the neuroendocrine cells of the gastrointestinal mucosa, the intestinal enterochromaffin cells and the main cells, and the submucosal cells of the colon [9,10]. The widespread distribution of ADM in the gastrointestinal tract provides an anatomical basis for regulating gastrointestinal physiology and pathology. For example, it has been reported that over-expression of ADM in the stomach can inhibit gastric acid secretion [11]. In other studies, ADM can protect mucosal as an endothelial cell growth factor by promoting mucosal healing [12], and has anti-inflammatory effects in a mouse DSS-induced colitis model [13]. However, the relationship between ADM and gastric inflammation especially in *H. pylori*-associated gastritis is presently unknown.

In our study, we have, for the first time, demonstrated a pro-inflammation role of ADM in *H. pylori* infection and *H. pylori*-induced gastritis. ADM is increased in gastric mucosa of both patients and mice infected with *H. pylori*. *H. pylori* induces ADM production from gastric epithelial cells in a *cagA*-dependent manner via activating PI3K-AKT signaling pathway. ADM derived from gastric epithelial cells promotes IFN-γ-producing T cells via the phosphorylation of AKT and STAT3. Besides, ADM also induces macrophages to produce IL-12, which promotes IFN-γ-producing T-cell responses. *In vivo*, the increased ADM promotes inflammation and IFN-γ-producing T-cell responses in the stomach during *H. pylori* infection, which contributes to *H. pylori*-associated gastritis. These findings provide a novel regulatory network involving ADM within the gastric microenvironment and a potentially new target of ADM in the treatment of *H. pylori*-associated gastritis.

## Results

### Adrenomedullin is increased in gastric mucosa of *H. pylori*-infected patients and mice

To evaluate the potential role of adrenomedullin (ADM) in *H. pylori*-associated pathology, first we compared the ADM levels in gastric tissues. Notably, compared to uninfected donors, the overall level of ADM mRNA was higher respectively in gastric mucosa of *H. pylori*-infected patients (Fig 1A). Next, ADM expression was positively correlated with *H. pylori* colonization (Fig 1B), suggesting induction of ADM by *H. pylori*.

**Fig 1.**
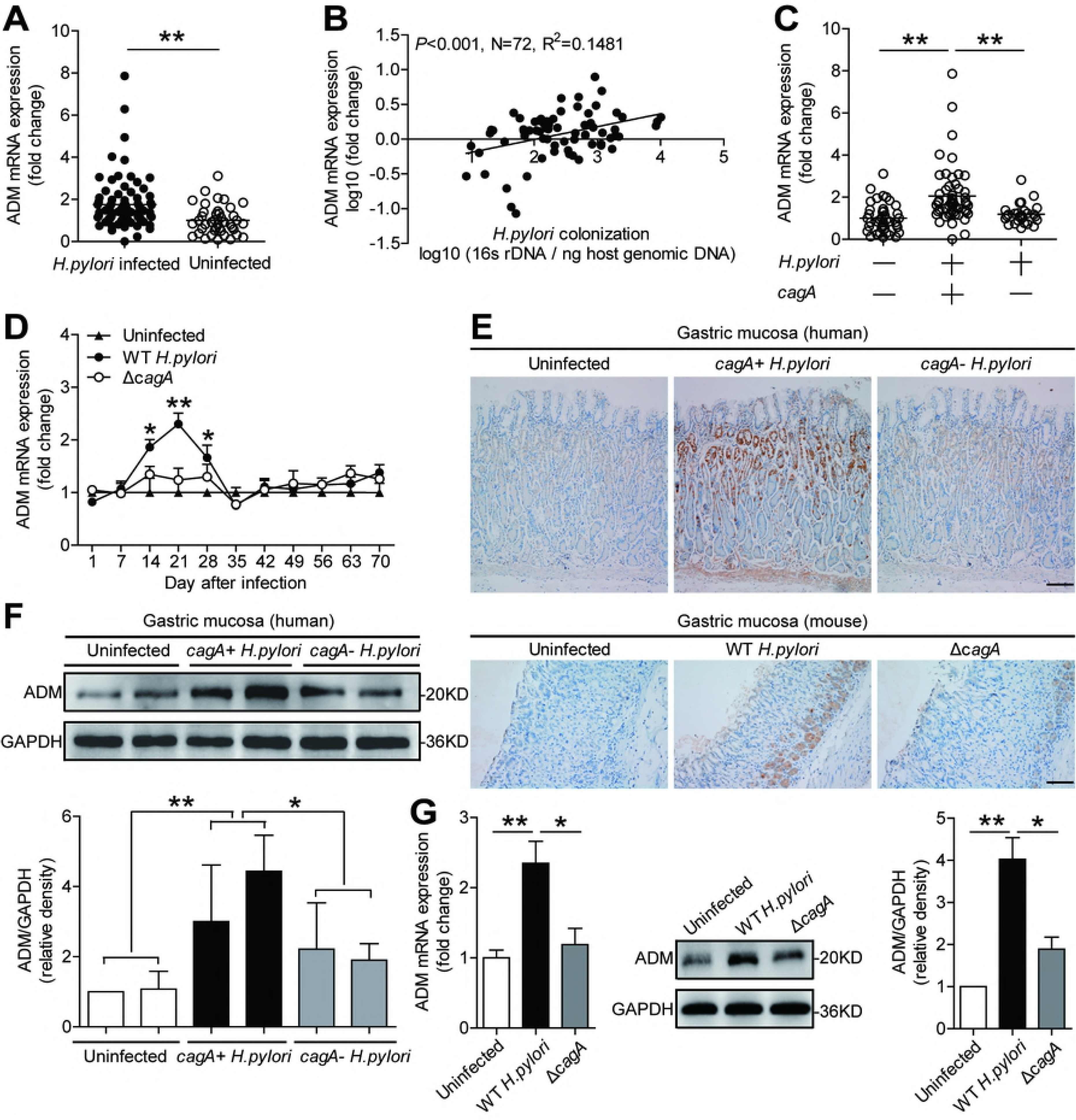
ADM is increased in gastric mucosa of *H. pylori*-infected patients and mice. (A) ADM mRNA expression in gastric mucosa of uninfected donors (n=41) and *H. pylori*-infected patients (n=80) was compared. (B) The correlation between ADM expression and *H. pylori* colonization in gastric mucosa of *H. pylori*-infected patients was analyzed. (C) ADM mRNA expression in gastric mucosa of uninfected donors (n=41), *cagA*^-^ *H. pylori*-infected (n=27), and *cagA*+ *H. pylori*-infected (n=53) patients was compared. (D) Dynamic changes of ADM mRNA expression in WT *H. pylori*-infected, *ΔcagA*-infected, and uninfected mice. n=5 per group per time point in D. (E) ADM protein in gastric mucosa of uninfected donors, *cagA*^-^ *H. pylori*-infected, and *cagA*+ *H. pylori*-infected patients or in gastric mucosa of WT *H. pylori*-infected, *ΔcagA*-infected, and uninfected mice on day 21 p.i. was analyzed by immunohistochemical staining. Scale bars: 100 microns. (F) ADM protein in gastric mucosa of uninfected donors, *cagA*^-^ *H. pylori*-infected, and *cagA*+ *H. pylori*-infected patients was analyzed by western blot and statistically analyzed (n=3). (G) ADM mRNA expression and ADM protein in human primary gastric mucosa from uninfected donors infected with WT *H. pylori* or *ΔcagA ex vivo* analyzed by real-time PCR and western blot and statistically analyzed (n=3). Results are representative of 3 independent experiments. The horizontal bars in panels A and C represent mean values. Each ring or dot in panels A, B and C represents 1 patient or donor. *P<0.05, and **P<0.01 for groups connected by horizontal lines compared, or compared with uninfected mice.

The presence of *cagA* is strongly associated with the development of *H. pylori*-associated gastritis [14]. Notably, we found that ADM mRNA expression in *cagA*-positive patients was significantly higher than that in *cagA*-negative individuals (Fig 1C). Immunohistochemical staining (Fig 1E) and western blot analysis (Fig 1F) showed that ADM protein in *cagA*-positive patients was also significantly higher than that in *cagA*-negative individuals. Consistent with our findings in humans, the levels of ADM mRNA (Fig 1D) and protein (Fig 1E) were almost only detected in WT *H. pylori*-infected mice, reaching a peak 21 days post infection (p.i.), indicating a role for *cagA* is involved in the induction of ADM in gastric mucosa during *H. pylori* infection. Finally, we examined expression of ADM in human primary gastric mucosa stimulated with *H. pylori*, and found that infection with WT *H. pylori* infection, the levels of ADM mRNA and protein in human primary gastric mucosa were significantly increased compared to the samples either not infected or infected with *ΔcagA* (Fig 1G). Taken together, these findings suggest that ADM is increased in *H. pylori*-infected gastric mucosa of patients and mice.

### Gastric epithelial cells stimulated by *H. pylori* express adrenomedullin

Gastric epithelial cells are known to be the first-contacting cell type in gastric mucosa during *H. pylori* infection [15]. We therefore sought to determine whether gastric epithelial cells were responsible for ADM expression during *H. pylori* infection. Interestingly, within gastric mucosa of *H. pylori*-infected donors, ADM was not only expressed in CD326^+^ gastric epithelial cells (Fig 2A and S1A Fig); but also expressed in H^+^/K^+^ ATPase^+^ parietal cells (Fig 2B and S1A Fig) and pepsinogen II^+^ chief cells (Fig 2C and S1A Fig). These data suggest that gastric epithelial cells are the source cells that express ADM in gastric mucosa during *H. pylori* infection.

**Fig 2.**
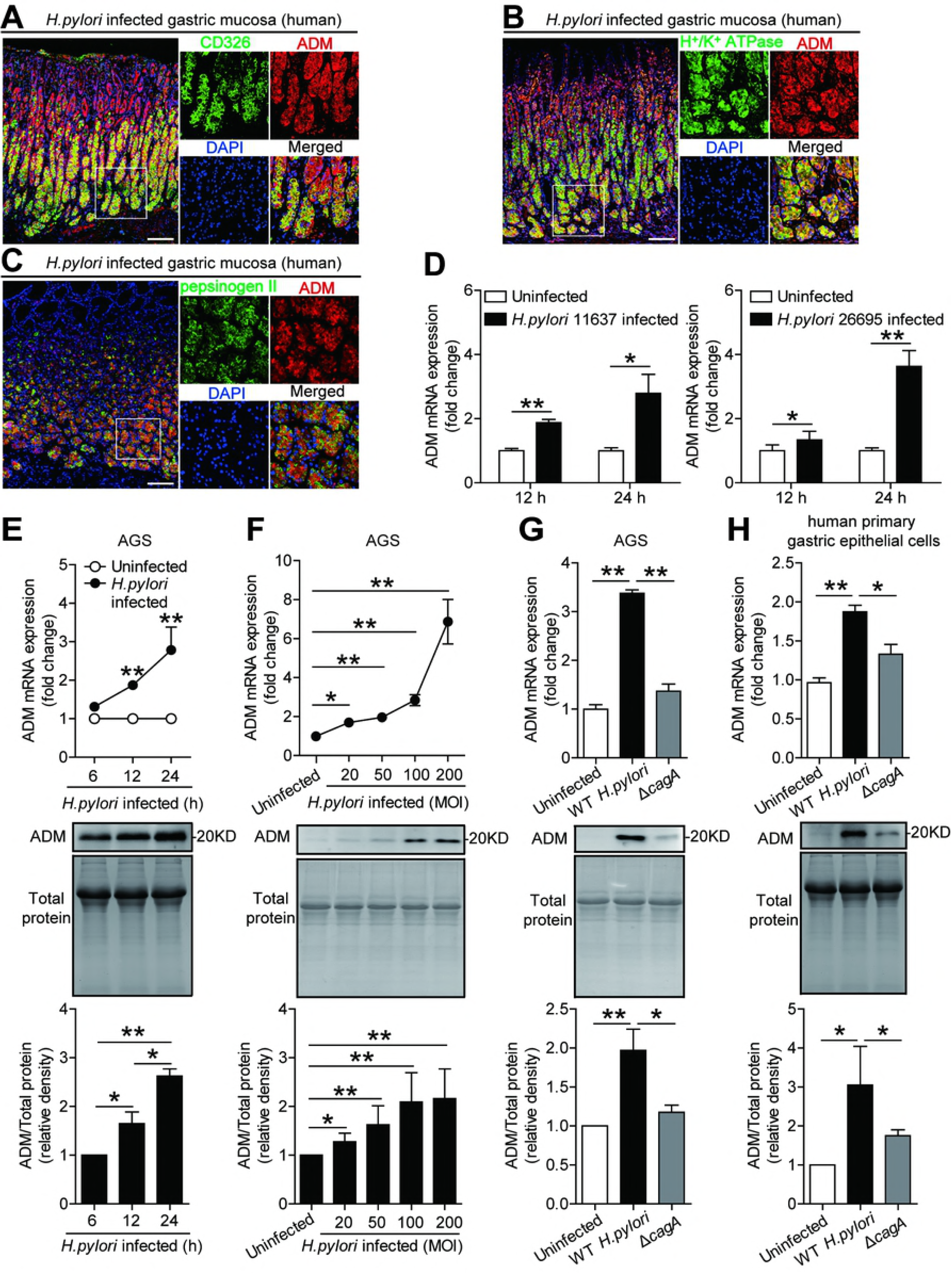
Gastric epithelial cells stimulated by *H. pylori* express ADM. (A) Representative immunofluorescence staining images showing ADM-expressing (red) CD326^+^ gastric epithelial cells (green) in gastric mucosa of *H. pylori*-infected patients. Scale bars: 100 microns. (B) Representative immunofluorescence staining images showing ADM-expressing (red) H+/K+ATPase+ parietal cells (green) in gastric mucosa of *H. pylori*-infected patients. Scale bars: 100 microns. (C) Representative immunofluorescence staining images showing ADM-expressing (red) pepsinogen II+ chief cells (green) in gastric mucosa of *H. pylori*-infected patients. Scale bars: 100 microns. (D) ADM mRNA expression in *H. pylori* 11637-infected, *H. pylori* 26695-infected and uninfected AGS cells at 12 or 24 h (MOI=100) was analyzed by real-time PCR (n=3). (E and F) ADM mRNA expression and ADM protein in/from WT *H. pylori*-infected and uninfected AGS cells at different time point (MOI=100) (E) or with different MOI (24 h) (F) were analyzed by real-time PCR and western blot and statistically analyzed (n=3). Results are representative of 3 independent experiments. Protein loading is shown in the coomassie brilliant blue staining gel. (G and H) ADM mRNA expression and ADM protein in/from WT *H. pylori*-infected, *ΔcagA*-infected, and uninfected AGS cells (G) or human primary gastric epithelial cells (H) (MOI=100, 24 h) were analyzed by real-time PCR and western blot and statistically analyzed (n=3). Results are representative of 3 independent experiments. Protein loading is shown in the coomassie brilliant blue staining gel. *P<0.05, and **P<0.01 for groups connected by horizontal lines compared, or compared with uninfected cells.

Next, we used two strains of *H. pylori* to infect AGS cells, an immortalized human gastric epithelial cell line, and found that *H. pylori* significantly increased ADM expression (Fig 2D). Moreover, *H. pylori*-infected AGS cells could increase ADM mRNA expression and ADM protein production in a time-dependent (Fig 2E) and infection dose-dependent manner (Fig 2F). Notably, compared to uninfected or the ones infected with *ΔcagA*, WT *H. pylori*-infected AGS cells (Fig 2G) and human primary gastric epithelial cells (Fig 2H) also potently increased ADM mRNA expression and ADM protein production. Similar observations were made when another human gastric epithelial cell line HGC-27 cells (S1B and C Fig). Collectively, these results demonstrate that *H. pylori* infection induces ADM expression in gastric epithelial cells, implying that induction of ADM in these cells are a major cause of increased ADM within the *H. pylori*-infected gastric mucosa.

### *H. pylori* stimulates gastric epithelial cells to express adrenomedullin via PI3K-AKT pathway

To see which signaling pathways might operate in the induction of ADM in gastric epithelial cells, we first per-treated AGS cells with corresponding inhibitors, and then stimulated AGS cells with *H. pylori*. We found that only blocking PI3K-AKT pathway with Wortmannin effectively suppressed ADM mRNA expression and ADM protein production in/from *H. pylori*-infected gastric epithelial cells (Fig 3A and B and S2 Fig). Furthermore, AKT, a direct PI3K-AKT pathway downstream substrate, was predominantly phosphorylated in AGS cells after stimulated with *H. pylori*, and this was more noticeable when infected with a WT *H. pylori* compared to *ΔcagA* (Fig 3C), and this phosphorylation was abolished when PI3K-AKT signal transduction pathway was blocked with inhibitor Wortmannin (Fig 3C). Furthermore, *H. pylori* induced AKT phosphorylation in AGS cells in a dose-dependent (Fig 3D) as well as in a time-dependent manners (Fig 3E). These data imply that activation of PI3K-AKT signaling pathway is crucial for ADM induction by *H. pylori* in gastric epithelial cells.

**Fig 3.**
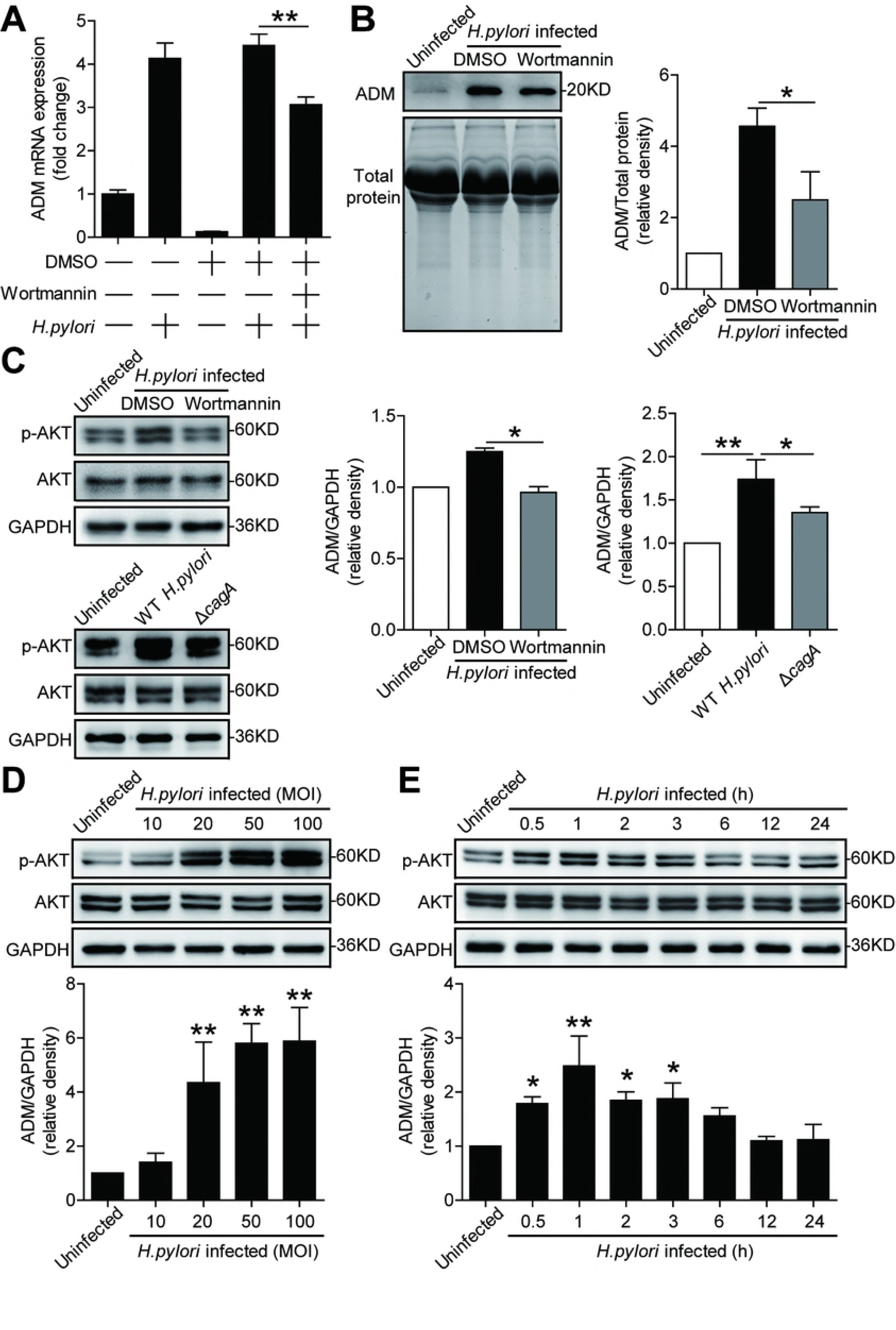
*H. pylori* stimulates gastric epithelial cells to express ADM via PI3K-AKT pathway. (A and B) AGS cells were pre-treated with Wortmannin (a PI3K-AKT inhibitor) and then stimulated with WT *H. pylori* (MOI=100) for 24 h. ADM mRNA expression (A) and ADM protein (B) in/from AGS cells were analyzed by real-time PCR and western blot and statistically analyzed (n=3). Results are representative of 3 independent experiments. (C) AGS cells were pre-treated with Wortmannin (a PI3K-AKT inhibitor) and then stimulated with WT *H. pylori* (MOI=100) for 1 h, or were infected with WT *H. pylori* or *ΔcagA* (MOI=100) for 1 h. AKT and p-AKT proteins were analyzed by western blot and statistically analyzed (n=3). Results are representative of 3 independent experiments. (D and E) AGS cells were infected with WT *H. pylori* with different MOI (1 h) (D) or at different time point (MOI=100) (E). AKT and p-AKT proteins were analyzed by western blot and statistically analyzed (n=3). Results are representative of 3 independent experiments. *P<0.05, and **P<0.01 for groups connected by horizontal lines compared.

### *In vivo* blockade of adrenomedullin significantly reduced inflammation and IFN-γ-producing T-cell responses in the stomach during *H. pylori* infection

To understand the possible biological effects of ADM induction during *H. pylori* infection, we compared ADM expression within the gastric mucosa with the severity of gastritis observed in patients infected with *H. pylori*. Notably, higher ADM expression was strongly associated with more severe gastritis (Fig 4A). This led us to hypothesize that ADM might exert pro-inflammatory effects during *H. pylori* infection and thus contribute to gastritis.

**Fig 4.**
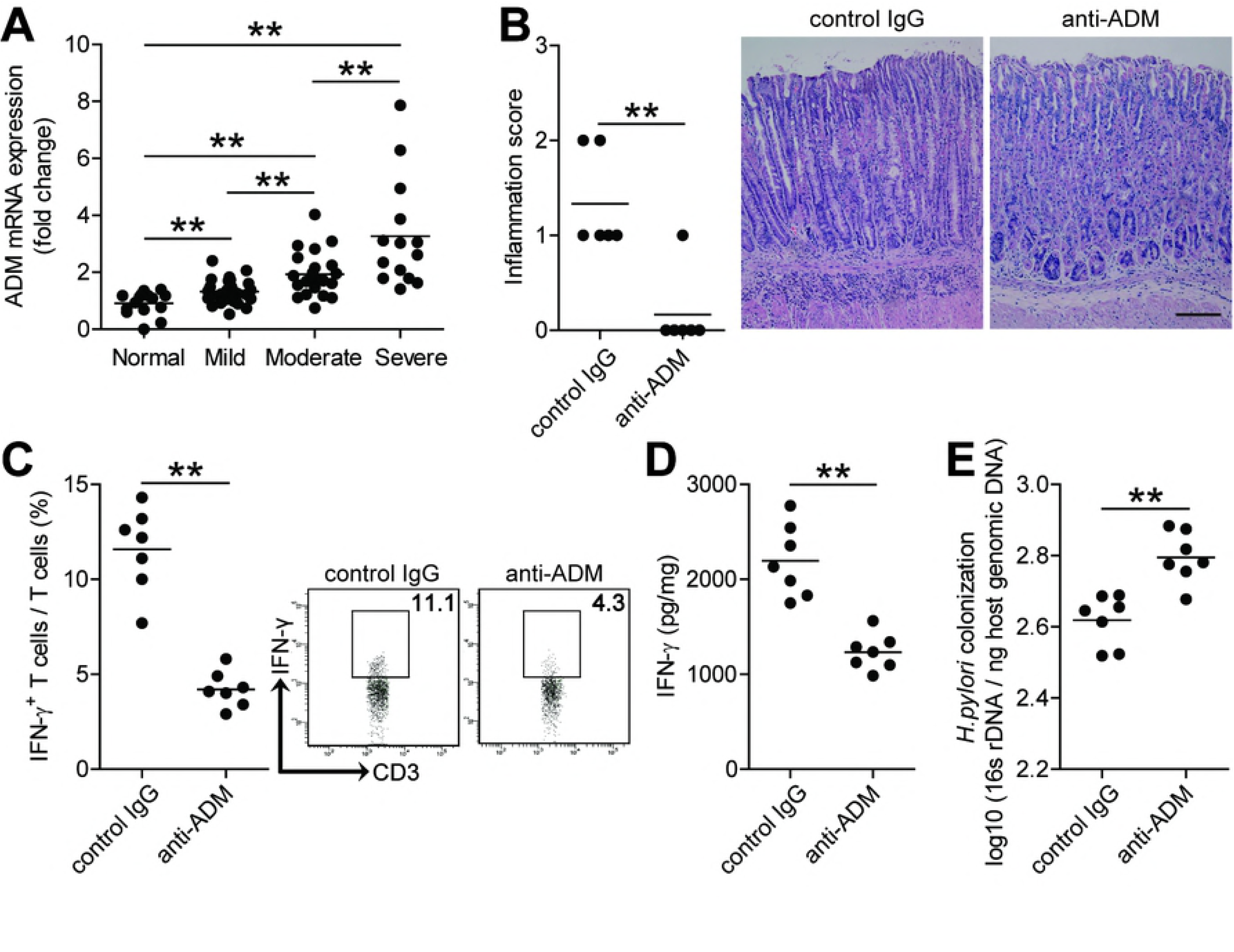
*In vivo* blockade of ADM significantly reduced inflammation and IFN-γ-producing T-cell responses in the stomach during *H. pylori* infection. (A) ADM mRNA expression in gastric mucosa of *H. pylori*-infected patients with normal gastric histopathology (n=32), or with mild (n=19), moderate (n=16), severe inflammation (n=12) was compared. (B) Histological scores of inflammation in gastric antra of WT *H. pylori*-infected mice injected with Abs against ADM or corresponding control IgG on day 21 p.i. were compared. H&E staining, scale bars: 100 microns. (C) The IFN-γ-producing T-cell responses in gastric mucosa of WT *H. pylori*-infected mice injected with Abs against ADM or corresponding control IgG on day 21 p.i. were compared. (D) The IFN-γ production in gastric mucosa of WT *H. pylori*-infected mice injected with Abs against ADM or corresponding control IgG on day 21 p.i. was compared. (E) The bacterial colonization in gastric mucosa of WT *H. pylori*-infected mice injected with Abs against ADM or corresponding control IgG on day 21 p.i. was compared. The horizontal bars in panels A, B, C, D and E represent mean values. Each dot in panels A, B, C, D and E represents 1 patient or mouse. **P<0.01 for groups connected by horizontal lines compared.

To test this hypothesis *in vivo*, we conducted a series of loss-of-function experiments involving ADM, and evaluated the inflammatory response in gastric mucosa on day 21 p.i.. Compared with mice treated with control IgG, mice treated with neutralizing antibodies against ADM showed significantly less inflammation in gastric mucosa (Fig 4B). Furthermore, neutralization of ADM significantly reduced the level of IFN-γ-producing T-cell responses (Fig 4C) and IFN-γ production (Fig 4D) in gastric mucosa but had no impact on Ly6G^-^CD11+monocytes, Ly6G+CD11b+ neutrophils, CD19+ B cells, and NK1.1+ natural killer cells (NK cells), IL-4-producing T cells and IL-17-producing T cells in gastric mucosa (S3 Fig). As for the control of bacteria growth by inflammatory immune response [16], finally, we compared the levels of bacterial colonization in gastric mucosa on day 21 p.i. and found that neutralization of ADM effectively increased *H. pylori* colonization (Fig 4E). Collectively, these results suggest that ADM has pro-inflammatory effects during *H. pylori* infection *in vivo* probably by regulating IFN-γ-producing T-cell responses.

### Adrenomedullin promotes IFN-γ-producing T-cell responses via PI3K-AKT and STAT3 activation

Next, we were therefore interested to know if and how ADM induces IFN-γ-producing T-cell responses. To begin, we found that an IFN-γ-producing T-cell infiltration as well as the expression of ADM receptor domain protein, receptor-modifying protein 2 (RAMP2), merged with CD3 staining on T cells in *H. pylori*-infected gastric mucosa (Fig 5A and S4A Fig). Moreover, an increased IFN-γ expression and a significant positive correlation between ADM and IFN-γ expression was found in *H. pylori*-infected gastric mucosa (Fig 5B and S4B Fig), altogether suggesting that T cells are a major target of ADM action within the inflamed gastric mucosa during *H. pylori* infection.

**Fig 5.**
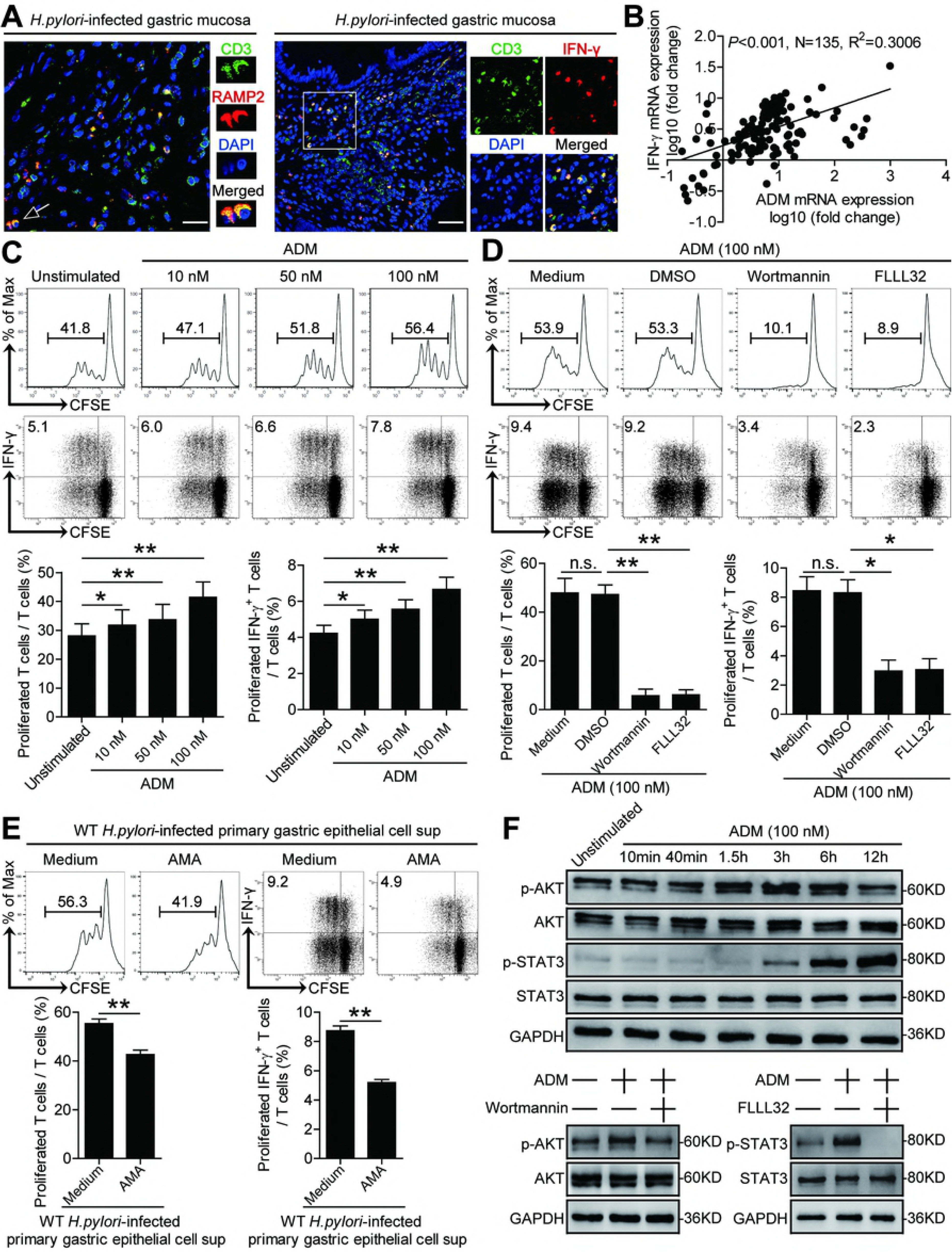
ADM promotes IFN-γ-producing T-cell responses via PI3K-AKT and STAT3 activation. (A) Representative immunofluorescence staining images showing RAMP2-expressing (red) CD3+ T cells (green) and IFN-γ-expressing (red) CD3+ T cells (green) in gastric mucosa of *H. pylori*-infected patients. Scale bars: 20 microns (left), 100 microns (right). (B) The correlation between ADM expression and IFN-γ expression in gastric mucosa of *H. pylori*-infected patients was analyzed. (C and D) T cells were stimulated with ADM (10, 50, 100 nM) (C), or pre-treated with Wortmannin (a PI3K-AKT inhibitor) and then stimulated with ADM (100 nM) (D) for 5 days as described in Methods. T cell proliferation and IFN-γ production was assessed by flow cytometry and statistically analyzed (n=3). Results are representative of 3 independent experiments. (E) T cells were stimulated with the culture supernatants from WT *H. pylori*-infected human primary gastric epithelial cells in the presence or absence of ADM fragment 22-52 (an ADM receptor antagonist) (AMA) for 5 days as described in Methods. T cell proliferation and IFN-γ production was assessed by flow cytometry and statistically analyzed (n=3). Results are representative of 3 independent experiments. (F) T cells were stimulated with ADM (100 nM) at different time point, or pre-treated with Wortmannin (a PI3K-AKT inhibitor) or FLLL32 (an STAT3 phosphorylation inhibitor) and then stimulated with ADM (100 nM) for 6 h. AKT and p-AKT proteins and STAT3 and p-STAT3 proteins were analyzed by western blot. *P<0.05, and **P<0.01 for groups connected by horizontal lines compared. sup, supernatants.

To further evaluate the contribution of ADM to the induction of IFN-γ-producing T-cell responses, we stimulated T cells with ADM and found that ADM was able to potently induce T cell proliferation and IFN-γ production in a dose-dependent manner (Fig 5C and S4D Fig). To see which signaling pathways might operate in the induction of IFN-γ-producing T cells by ADM, we first per-treated T cells with corresponding inhibitors, and then stimulated T cells with ADM. We found that blocking PI3K-AKT pathway with Wortmannin or STAT3 phosphorylation with FLLL32 effectively suppressed T cell proliferation and IFN-γ production (Fig 5D and S4C and E Fig). Furthermore, AKT and STAT3 were predominantly phosphorylated in T cells after stimulated with ADM in a time-dependent manner (Fig 5F and S4F Fig), and this phosphorylation was abolished when PI3K-AKT signal transduction pathway was blocked with Wortmannin or STAT3 phosphorylation was abolished with FLLL32 (Fig 5F and S4F Fig). Finally, to ascertain whether ADM from *H. pylori*-stimulated gastric epithelial cells contribute to the induction of IFN-γ-producing T-cell responses, we abolished ADM-ADM receptor interaction with ADM fragment 22-52 (an ADM receptor antagonist) (AMA) on T cells, and then stimulated T cells with the culture supernatants from WT *H. pylori*-stimulated primary gastric epithelial cells. As expected, blocking ADM-ADM receptor interaction effectively inhibited T cell proliferation and IFN-γ production (Fig 5E). Taken together, these findings suggest that ADM promotes IFN-γ-producing T-cell responses via activation of PI3K-AKT and STAT3 signaling pathways.

### Adrenomedullin-stimulated macrophages induce IFN-γ-producing T-cell responses

It has previously been shown that IFN-γ-producing T cells are induced by IL-12 [17], and that mascrophages are potent producers of IL-12 at sites of bacterial infection [18]. Since macrophages have previously been implicated in *H. pylori* gastritis [19], and we next were interested to learn whether ADM-regulated macrophages would affect IFN-γ-producing T-cell responses via IL-12 during *H. pylori* infection. To begin, we found that an expression of ADM receptor domain protein, RAMP2, merged with CD68 staining on macrophages in *H. pylori*-infected gastric mucosa (Fig 6A and S5A Fig). Next, ADM was able to potently induce IL-12 expression and production from macrophages (Fig 6B). Moreover, an increased IL-12 expression and significant positive correlations between ADM and IFN-γ expression or between IFN-γ and IL-12 expression were found in *H. pylori*-infected gastric mucosa (Fig 6C and S5AB Fig), Taken together, these findings suggesting that macrophages are another major target of ADM action and may be responsible for the IFN-γ-producing T-cell responses within the inflamed gastric mucosa during *H. pylori* infection.

**Fig 6.**
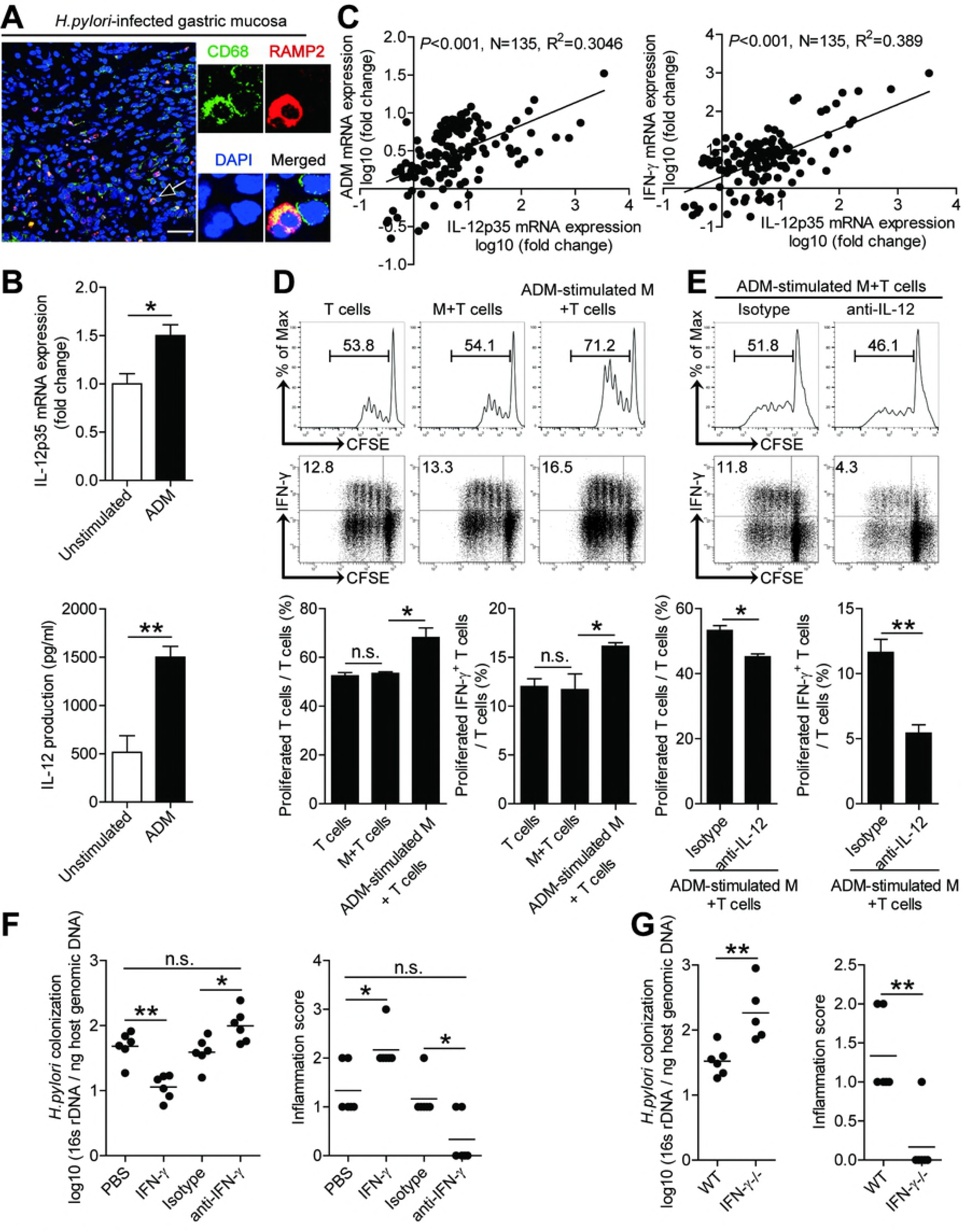
ADM-stimulated macrophages induce IFN-γ-producing T-cell responses. (A) Representative immunofluorescence staining images showing RAMP2-expressing (red) CD68^+^ macrophages (green) in gastric mucosa of *H. pylori*-infected patients. Scale bars: 50 microns. (B) IL-12p35 mRNA expression and IL-12 protein in/from unstimulated or ADM-stimulated human primary macrophages derived from blood monocytes were analyzed by real-time PCR and ELISA (n=3). (C) The correlations between ADM expression and IL-12p35 expression or IL-12p35 expression and IFN-γ expression in gastric mucosa of *H. pylori*-infected patients were analyzed. (D and E) T cell proliferation and IFN-γ production of T cell-macrophage co-culture was assessed by flow cytometry as described in Methods and statistically analyzed (n=3). (F) Histological scores of inflammation and bacteria colonization in gastric mucosa of WT *H. pylori*-infected mice injected with IFN-γ or PBS control, Abs against IFN-γ or corresponding control isotype IgG on day 21 p.i. were compared. (G) Histological scores of inflammation and bacteria colonization in gastric mucosa of WT *H. pylori*-infected WT and IFN-γ^-/-^ mice on day 21 p.i. were compared. The horizontal bars in panels F and G represent mean values. Each dot in panels C, F and G represents 1 patient or mouse. *P<0.05, **P<0.01, and n.s. P>0.05 for groups connected by horizontal lines compared.

To further evaluate the contribution of ADM-regulated macrophages to the induction of IFN-γ-producing T-cell responses, we stimulated macrophages with ADM and then co-cultured with T cells, and found that ADM-stimulated macrophages was able to potently induce T cell proliferation and IFN-γ production (Fig 6D and S5C Fig). To see whether IL-12 from ADM-stimulated macrophages might operate in the induction of IFN-γ-producing T cells, we added with neutralizing antibodies against IL-12 in co-culture system above, and found that blocking IL-12 effectively suppressed T cell proliferation and IFN-γ production induced by ADM-stimulated macrophages (Fig. 6E and S5DA Fig). Taken together, these findings suggest that ADM-stimulated macrophages promote IFN-γ-producing T-cell responses via IL-12.

Finally, we conducted a series of loss- and gain-of-function *in vivo* experiments involving IFN-γ, and evaluated the inflammatory response and bacterial colonization in gastric mucosa on day 21 p.i.. Compared with mice treated with control isotype IgG, mice treated with neutralizing antibodies against IFN-γ showed significantly less inflammation and higher *H. pylori* colonization in gastric mucosa (Fig 6F). Conversely, injection of IFN-γ significantly increased inflammation and reduced *H. pylori* colonization in gastric mucosa (Fig 6F). Similarly, compared to WT mice, IFN-γ^-/-^ mice showed significantly less inflammation and higher *H. pylori* colonization in gastric mucosa (Fig 6G). Collectively, these results suggest that ADM-stimulated macrophages promote IFN-γ-producing T-cell responses and IFN-γ contributes to inflammation during *H. pylori* infection.

## Discussion

More than 50% of the world’s population infects *H. pylori* in their upper gastrointestinal tracts, which is more common in developing countries than Western countries [1]. *H. pylori* infection is a threat to human health, for example, the long-term colonization of *H. pylori* in the stomach can change PH of the stomach, promote chronic gastritis, gastric ulcers, even gastric cancer [2]. However, until now, the mechanism of *H. pylori*-associated chronic gastritis remains unclear, and it is believed that the interplays between host and bacterial virulence factors [14,20], such as VacA [21], urease [22], especially a major virulence factor *cagA* that can be injected into host cell by Type IV Secretion System (T4SS) [23], and the following persistent inflammation are likely the underlying causes.

In this study we demonstrated a multistep model of inflammation during *H. pylori* infection within the gastric mucosa involving interactions among *H. pylori*, gastric epithelial cells, T cells and macrophages via ADM (Fig 7). *In vivo* and *in vitro*, we established that *H. pylori*-associated virulence factor *cagA* was essential to inducing ADM expression in gastric epithelial cells, which in turn promoted gastric inflammation, CD3+T cell proliferation and IFN-γ production. Moreover, the increased ADM induced macrophages to secrete IL-12, which also promoted IFN-γ-producing T-cell responses. To our knowledge, this study is the first time to demonstrate the pro-inflammatory role of ADM and its association with macrophages and T cells in *H. pylori*-induced gastritis.

**Fig 7.**
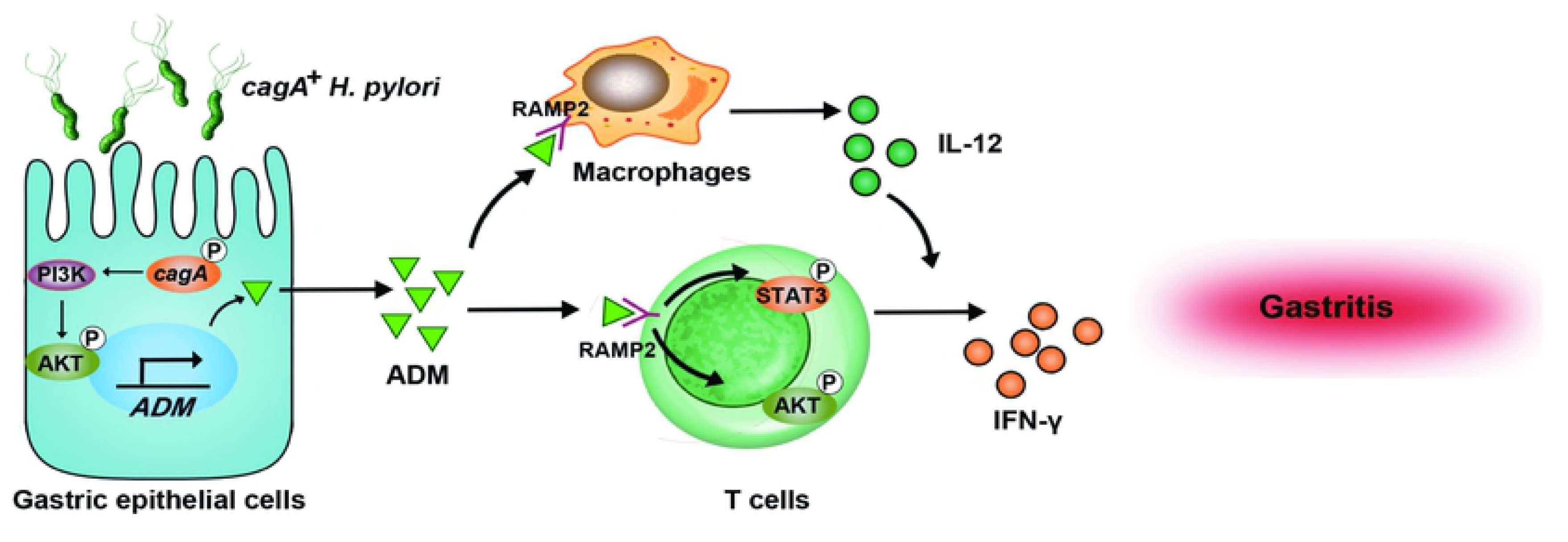
A proposed model of cross-talk among *H. pylori*, gastric epithelial cells, ADM, and macrophages leading to IFN-γ-producing T-cell responses and ADM-IFN-γ-mediated pro-inflammation in gastric mucosa during *H. pylori* infection. *H. pylori* stimulates gastric epithelial cells to secrete ADM via PI3K-AKT signaling pathway activation in a *cagA*-dependent manner. Release of ADM induces the differentiation of IFN-γ-producing T cells via the phosphorylation of AKT and STAT3. On the other hand, ADM stimulates macrophages to produce IL-12, which promotes the IFN-γ-producing T-cell responses. Increased ADM and IFN-γ exert pro-inflammatory effect within the gastric microenvironment which collectively contributes to gastritis during *H. pylori* infection.

ADM is a biologically active peptide which was isolated in 1993 by Kitamura et al [24] from human pheochromocytoma. It was discovered that ADM was able to activate platelet adenylate cyclase and exerted a long-lasting hypotensive effect in rats [24]. The expression of ADM is related to many factors, for example, inflammatory factors IL-1β and TNF-α can induce ADM expression in some cell types [25]. Some studies showed that plasmatic ADM levels were raised in many infectious diseases, particularly in sepsis [26], suggesting a inflammatory role of this hormone. In regard to the digestive system, several studies have demonstrated that ADM levels rise in infected gastrointestinal tissues. For example, examination of ADM expression in gastrointestinal epithelium revealed higher mRNA levels in paratuberculosis-infected versus uninfected cows [27]. It has been reported that infection of *H. pylori* would upregulate the expression of ADM in gastric epithelial cells *in vitro* [28]. In our study, firstly we validated the previous results [28] *in vitro*, besides, we confirmed the increased expression of ADM in gastric tissues from 135 *H. pylori*-infected patients as well as in the mouse models infected with *H. pylori in vivo*.

ADM can activate a great variety of signal transduction pathways, depending on the species, organ, tissue, and cell type. However, there are three main signaling pathways by which ADM exerts its actions: cAMP, phosphoinositide 3-kinase (PI3K)-AKT, and mitogen-activated protein kinase (MAPK)-extracellular signal-regulated protein kinase (ERK) [24,29,30]. All signal mechanisms in which ADM is involved are the basis of this peptide’s extensive repertoire of biological functions such as vasodilatation, cellular proliferation, apoptosis modulation, or in?ammatory regulation [30-33]. It has been reported that ADM can promote the proliferation of cells such as cardiomyocytes and vascular endothelial cells, via signaling pathways such as PI3K-AKT and MAPK by acting on ADM receptor 1 (ADMR1) and ADMR2, and participate in the construction of blood vessels. Simultaneously, studies have shown that ADM receptors are expressed on T cells [34]. However, the direct effect of ADM on T cells in the gastrointestinal tract has not been reported. Our results for the first time show that, during *H. pylori* infection, gastric epithelial cells-derived ADM can directly promote the proliferation of T cells and the production of IFN-γ from T cells through PI3K-AKT and STAT3 signaling pathways. The significance of this finding is to further understand the role of ADM in the alteration of the immune environment in the *H. pylori*-infected stomach.

In our study, we found that after blocking ADM during *H. pylori* infection, gastric inflammation decreased *in vivo*. However, some studies reported that ADM showed potent anti-inflammatory activity by down-regulating the production of a wide panel of inflammatory mediators such as IL-10 and TNF-α by microglia and DCs [35]. We guess that ADM may exert different roles by acting on different types of target cells in different diseases. Here, we found that ADM from *H. pylori*-infected gastric epithelial cells directly promoted T cell proliferation and IFN-γ production. Besides, we also found that ADM indirectly acted on T cells by IL-12 derived by ADM stimulated-macrophages, which resembles the data on IL-1β secretion in synovial inflammation [36]. It is clear that a mixed acute and chronic inflammatory reaction takes place in *H. pylori*-induced inflammatory environments, where multifarious immune cell types infiltrate and play complicated roles [37,38]. In a recent report, TLR9^-/-^ mice were found to show increased signs of gastritis upon *H. pylori* infection [39]. This report supports our findings that ADM may play a role in promoting inflammation by directly or indirectly acting on immune cells during *H. pylori* infection.

To sum up, our findings provide a novel regulatory network involving *H. pylori*, gastric epithelial cells, ADM, macrophages, T cells, and IFN-γ in *H. pylori*-induced gastritis. Given the notable relationship between the level of ADM and the severity of gastric inflammation observed in *H. pylori*-infected patients, it is possible that ADM might serve as a novel diagnostic and prognostic biomarker for *H. pylori-*associated gastritis. Future clinical studies are necessary to investigate and verify the ADM-associated mechanisms in humans, which may lead to the application of novel pharmacologic approaches to resist this gastric inflammation.

## Materials and methods

### Patients and specimens

The gastric biopsy specimens and blood were collected from 135 *H. pylori*-infected and 41 uninfected patients who underwent upper esophagogastroduodenoscopy for dyspeptic symptoms at XinQiao Hospital (Supplementary Table 1). *H. pylori* infection was determined by [^14^C] urea breath test and rapid urease test of biopsy specimens taken from the antrum, and subsequently conformed by real-time PCR for 16s rDNA and serology test for specific anti-*H. pylori* antibodies (Abs). For isolation of human primary gastric epithelial cells, fresh non-tumor gastric tissues (at least 5-cm distant from the tumor site) were obtained from gastric cancer patients who underwent surgical resection and were determined as *H. pylori*-negative individuals as above at the Southwest Hospital. None of these patients had received chemotherapy or radiotherapy before sampling. Individuals with atrophic gastritis, hypochlorhydria, antibiotics treatment, autoimmune disease, infectious diseases and multi-primary cancer were excluded. The study was approved by the Ethics Committee of XinQiao Hospital and Southwest Hospital of Third Military Medical University. The written informed consent was obtained from each subject.

### Mice

All breeding and experiments were undertaken with review and approval from the Animal Ethical and Experimental Committee of Third Military Medical University. C57BL/6 interferon-γ^-/-^ (IFN-γ^-/-^) mice were kindly provided by Dr. Richard A. Flavell (Yale University). All mice used in experiments were female and viral Ab free for pathogenic murine viruses, negative for pathogenic bacteria including *Helicobacter* spp. and parasites, and were maintained under specific pathogen free (SPF) conditions in a barrier sustained facility and provided with sterile food and water.

### Antibodies and other reagents

Details are available in Supplementary Table 2.

### Bacterial culture and infection of mice with bacteria

*H. pylori* NCTC 11637 (*cagA* positive) (WT *H. pylori*) and *cagA*-knockout mutant *H. pylori* NCTC 11637 (*ΔcagA*) were grown in brain-heart infusion plates containing 10% rabbit blood at 37°C under microaerophilic conditions. For infecting mouse, bacteria were propagated in Brucella broth with 5 % fetal bovine serum (FBS) with gentle shaking at 37°C under microaerobic conditions. After culture for 1 d, live bacteria were collected and adjusted to 10^9^ CFU/ml. The mice were fasted overnight and orogastrically inoculated twice at a 1 d interval with 3×10^8^ CFU bacteria. Age-matched control mice were mock-inoculated with Brucella broth. Five to seven mice per group per time point were used for the experiments. *H. pylori* infection status and *H. pylori*-induced gastritis in murine experiments were confirmed using real-time PCR of *H. pylori* 16s rDNA, urease biopsy assays, Warthin-Starry staining and immunohistochemical staining for *H. pylori*, and evaluation of inflammation by haematoxylin and eosin (H&E) staining (data not shown).

### Preparation of rabbit-anti-mouse ADM Ab

Polyclone Abs against ADM were produced by ChinaPeptides Co., Ltd. Female New Zealand rabbits received injections at multiple subcutaneous sites with 100 ug of purified peptide emulsified with complete Freund’s adjuvant according to their protocols. Then the rabbits were further immunized at 2-week intervals with 100 ug ADM emulsified with incomplete Freund’s adjuvant. The anti-sera obtained after the fourth booster injection were screened for anti-ADM activity, and then affinity purified on CNBR-activated sepharose 4b Fast Flow columns.

### *In vivo* blockade of ADM

One hour before infection with WT *H. pylori*, rabbit-anti-mouse ADM Ab (20 ug), or normal rabbit IgG (20 ug) was administered to each mouse via intraperitoneal injection, and the administration was repeated every week until the mice were sacrificed at the indicated time.

### *In vivo* blockade of IFN-γ or IFN-γ administration

One hour before infection with WT *H. pylori*, rabbit-anti-mouse IFN-γ mAb (20 ug), or control isotype IgG (20 ug); mouse IFN-γ (20 ug), or equal volume of sterile PBS was administered to each mouse via intraperitoneal injection, and the administration was repeated every week until the mice were sacrificed at the indicated time.

### Evaluation of inflammation and bacterial load

The mice were sacrificed at the indicated time. The stomach was cut open from the greater curvature and half of the tissue was cut into four parts for RNA, DNA, tissue fixation for H&E staining and protein extraction, respectively. The degree of inflammation was evaluated in a blinded fashion by two pathologists, and each section was given a score of 0–5 according to the established criteria [40]. DNA of the biopsy specimens were extracted with QIAamp DNA Mini Kit. The density of *H. pylori* colonization was quantified by real-time PCR, detecting *H. pylori*-specific 16s rDNA as previously described [41] using specific primer and probe (Supplementary Table 3). Expression of 16s rDNA was measured using the TaqMan method. The amount of mouse β2-microglobulin DNA in the same specimen was measured to normalize the data. The density of *H. pylori* in the samples was expressed as the number of bacterial genomes per nanogram of host genomic DNA according to a previous report [42]. Another half of stomach was used for isolation of single cells. The isolated single cells were collected and analyzed by flow cytometry.

### Isolation of single cells from tissues

Fresh tissues were washed three times with Hank’s solution containing 1% FBS, cut into small pieces, collected in RPMI 1640 containing 1 mg/ml collagenase IV and 10 mg/ml DNase I, and then mechanically dissociated by using the gentle MACS Dissociator (Miltenyi Biotec). Dissociated cells were further incubated for 0.5-1 h at 37°C under continuous rotation. The cell suspensions were then filtered through a 70-μm cell strainer (BD Labware).

### Human gastric epithelial cell/tissue culture and stimulation

Primary gastric epithelial cells were purified from gastric tissue single-cell suspensions from uninfected donors with a MACS column purification system using anti-human CD326 magnetic beads. The sorted primary gastric epithelial cells were used only when their viability was determined >90% and their purity was determined >95%. For human gastric epithelial cell lines (AGS cells and HGC-27 cells), 3×10^5^ cells per well in 12-well cell culture plate (for real-time PCR) or 1×10^6^ cells per well in 6-well cell culture plate (for western blot and ELISA) were starved in DMEM (Dulbecco’s Modified Eagle Medium)/F-12 medium supplemented with penicillin (100?U/ml) and streptomycin (100?μg/ml) for 6 h in a humidified environment containing 5% CO_2_ at 37?°C. Then the cells were incubated in antibiotic-free DMEM/F-12 medium supplemented with 10% FBS instead. The cell lines were used when their viability was determined >90%. Human gastric epithelial cell lines, primary gastric epithelial cells, or primary gastric mucosa tissues from uninfected donors were stimulated with WT *H. pylori* or *ΔcagA* at a multiplicity of infection (MOI) of 100 for 24 h. AGS cells were also stimulated with WT *H. pylori* at different MOI (24 h) or at the indicated time points (MOI=100). For signal pathway inhibition experiments, AGS cells were pre-treated with 5 μl (20 μM) SP600125 (a JNK inhibitor), SB203580 (a p38 MAPK inhibitor), AG490 (a JAK inhibitor), or Wortmannin (a PI3K-AKT inhibitor) for 2 h. After co-culture, cells were collected for real-time PCR, and western blot, and the culture supernatants were harvested for ELISA or were concentrated for western blot.

### T cell culture and stimulation

Human T cells were isolated from peripheral blood mononuclear cells (PBMCs) of uninfected donors, and cultured as described previously [43]. Briefly, in a 5-day incubation, bead-purified peripheral CD3^+^ T cells (2×10^5^ cells/well in 96-well plates) were labeled with carboxyfluorescein succinimidyl ester (CFSE) and were stimulated with various concentrations of ADM (10-100 nM) in 200 μl RPMI-1640 medium containing human recombinant (hr) IL-2 (20 IU/ml), anti-CD3 (2 μg/ml), and anti-CD28 (1 μg/ml) antibodies. For signal pathway inhibition experiments, T cells were pre-treated with 5 μl (20 μM) SP600125 (a JNK inhibitor), SB203580 (a p38 MAPK inhibitor), U0126 (an MEK-1 and MEK-2 inhibitor), Wortmannin (a PI3K-AKT inhibitor) or FLLL32 (an STAT3 phosphorylation inhibitor) for 2 h. In other cases, T cells were stimulated with the culture supernatants from human primary gastric epithelial cells infected with WT *H. pylori* (MOI=100, 24 h). In some cases, ADM fragment 22-52 (an ADM receptor antagonist) (1 μg/ml) was added into the culture supernatants and incubated for 2 h before stimulation. After 5-day co-culture, cells were collected for flow cytometry and western blot, and the supernatants were harvested for ELISA.

### Macrophage culture and stimulation

Human CD14^+^ cells were isolated from PBMCs of uninfected donors, and cultured as described previously.^43^To get macrophages, bead-purified CD14^+^ cells (2×10^4^ cells/well in 96-well plates) were stimulated with GM-CSF (100 ng/ml) for 5 days. Macrophages were stimulated with ADM (100 nM) for 24 h, and the cells were collected for real-time PCR, and the supernatants were harvested for ELISA. In other cases, after stimulated with ADM (100 nM) for 24 h, macrophages were then co-cultured with CFSE-labeled T cells (2×10^5^ cells/well in 96-well plates) in new complete RPMI-1640 medium containing rhIL-2 (20 IU/ml), anti-CD3 (2 μg/ml), and anti-CD28 (1 μg/ml) antibodies. In some cases, anti-human IL-12 Ab (20 μg/ml) or control IgG was added into the co-culture supernatants. After 5-day co-culture, T cells were collected for flow cytometry, and the supernatants were harvested for ELISA.

### Immunohistochemistry

Gastric tissue samples were fixed with paraformaldehyde and embed with paraffin, and then were cut into 5 µm sections. For immunohistochemical staining goat anti-mouse ADM Abs was used as primary Ab, and rabbit anti-goat-HRP as the second Ab, then sections were stained by DAB reagent. After that, all the sections were counterstained with hematoxylin and reviewed using a microscope (Nikon Eclipse 80i, Nikon).

### Immunofluorescence

Paraformaldehyde-fixed sections of gastric tissues were washed in PBS and blocked for 30 min with 20% goat serum in PBS and stained for ADM, CD326, H^+^/K^+^ ATPase, pepsinogen II, CD3, IFN-γ, CD68 and RAMP2. Slides were examined with a confocal fluorescence microscope (TCS SP8, Leica).

### Real-time PCR

DNA of the biopsy specimens were extracted with QIAamp DNA Mini Kit and RNA of biopsy specimens and cultured cells were extracted with TRIzol reagent. The RNA samples were reversed transcribed to cDNA with PrimeScriptTM RT reagent Kit. Real-time PCR was performed on the IQ5 (Bio-Rad) with the Real-time PCR Master Mix according to the manufacturer’s specifications. The mRNA expression of 16s rDNA, ADM, IFN-γ and IL-12p35 genes was measured using the TaqMan and/or SYBR green method with the relevant primers (Supplementary Table 3). For mice, mouse β2-microglobulin mRNA level served as a normalizer, and its level in the stomach of uninfected or WT mice served as a calibrator. For human, human glyceraldehyde 3-phosphate dehydrogenase (GAPDH) mRNA level served as a normalizer, and its level in the unstimulated cells or stomach of uninfected donors served as a calibrator. The relative gene expression was expressed as fold change of relevant mRNA calculated by the ΔΔCt method.

### Enzyme-linked immunosorbent assay (ELISA)

Human and mouse gastric tissues from specimens were collected, homogenized in 1 ml sterile Protein Extraction Reagent, and centrifuged. Tissue supernatants were collected for ELISA. Concentrations of IFN-γ in the tissue supernatants; concentrations of IFN-γ or IL-12 in T cell culture supernatants or macrophage culture supernatants were determined using ELISA kits according to the manufacturer’s instructions.

### Western blot analysis

Western blot assays were performed on 10%-15% SDS-PAGE gels transferred PDF membranes using equivalent amounts of cell or tissue lysate proteins of samples or the concentrated culture supernatants. Three percent BSA was used for blocking the PDF membranes. Human ADM was detected with rabbit anti-ADM Abs; human p-AKT, AKT, p-STAT3, STAT3 and GAPDH were detected with rabbit anti-p-AKT Abs, mouse anti-AKT Abs, rabbit anti-p-STAT3 Abs, rabbit anti-STAT3 Abs and mouse anti-GAPDH Abs, respectively. This was followed by incubation with HRP-conjugated secondary Abs. Bound proteins were visualized by using SuperSignal^®^ West Dura Extended Duration Substrate kit.

### Flow cytometry

Flow cytometric analysis was performed according to standard protocols. For intracellular cytokine measurements, the cells were stimulated with Leukocyte activation cooktail with BD GolgiPlug for 5 h. Intracellular cytokine staining was performed after fixation and permeabilization using Perm/Wash solution. The cells were analyzed by multicolor flow cytometry with FACSCanto II (BD Biosciences). Data were analyzed with FlowJo software (TreeStar) or FACSDiva software (BD Biosciences).

### Statistical analysis

Results are expressed as mean ± SEM. Student *t* test was generally used to analyze the differences between two groups, but when the variances differed, the Mann-Whitney U test was used. Inflammation score data were analyzed by the Mann-Whitney U test. Correlations between parameters were assessed using Pearson correlation analysis and linear regression analysis, as appropriate. SPSS statistical software (version 13.0) was used for all statistical analysis. All data were analyzed using two-tailed tests, and P<0.05 was considered statistically significant.

### Ethics statement

The study was approved by the Ethics Committee of XinQiao Hospital and Southwest Hospital of Third Military Medical University. All human subjects were adult, and the written informed consent was obtained from each subject. All animal experiments were performed in strict accordance with the Guide for the Care and Use of Laboratory Animals issued by the Ministry of Science and Technology of the People’s Republic of China. All breeding and experiments were undertaken with review and approval from the Animal Ethical and Experimental Committee of Third Military Medical University. The project license number is 201706A033.

## Acknowledgements

We wish to thank Yong-liang Zhao and Gang Guo for their helpful comments and constructive suggestions.

## Supporting information

**S1 Fig. *H. pylori* stimulates gastric epithelial cells express ADM.** (A) Secondary antibody-only stained sections as controls for the staining in Figure 2A-C. Scale bars: 100 microns. (B) ADM mRNA expression in *H. pylori* 11637-infected, *H. pylori* 26695-infected and uninfected HGC-27 cells at 12 or 24 h (MOI=100) was analyzed by real-time PCR (n=3). ADM mRNA expression and ADM protein in/from WT *H. pylori*-infected, *ΔcagA*-infected, and uninfected HGC-27 cells (MOI=100, 24 h) were analyzed by real-time PCR and western blot and statistically analyzed (n=3). *P<0.05, and **P<0.01 for groups connected by horizontal lines compared.

**S2 Fig. *H. pylori* stimulates gastric epithelial cells to express ADM via PI3K-AKT pathway.** AGS cells were pre-treated with signal pathway inhibitors and then stimulated with WT *H. pylori* (MOI=100) for 24 h. ADM mRNA expression in AGS cells was compared (n=3). **P<0.01, and n.s. P>0.05 for groups connected by horizontal lines compared.

**S3 Fig. *In vivo* blockade of ADM significantly reduced inflammation and IFN-γ-producing T-cell responses in the stomach during *H. pylori* infection.** The levels of Ly6G^-^CD11b+ monocytes, Ly6G+CD11b+ neutrophils, NK1.1+ natural killer cells (NK cells), CD19+ B cells, IL-4+ T cells, and IL-17+ T cells in gastric mucosa of WT *H. pylori*-infected mice injected with Abs against ADM or corresponding control IgG on day 21 p.i. were compared (n=5). n.s. P>0.05 for groups connected by horizontal lines compared.

**S4 Fig. ADM promotes IFN-γ-producing T-cell responses via PI3K-AKT and STAT3 activation.** (A) Secondary antibody-only stained sections as controls for the stainings in Figure 5A. Scale bars: 20 microns (left), 100 microns (right). (B) IFN-γ mRNA expression in gastric mucosa of uninfected donors (n=40) and *H. pylori*-infected patients (n=68) was compared. IFN-γ mRNA expression in gastric mucosa of uninfected donors (n=40), *cagA*^-^ *H. pylori*-infected (n=23), and *cagA*+*H. pylori*-infected (n=45) patients was compared. (C) T cells were pre-treated with signal pathway inhibitors and then stimulated with ADM (100 nM) for 5 days as described in Methods. T cell proliferation and IFN-γ production was assessed by flow cytometry. Results are representative of 3 independent experiments. (D and E) T cells were stimulated with ADM (10, 50, 100 nM) (D), or pre-treated with Wortmannin (a PI3K-AKT inhibitor) and then stimulated with ADM (100 nM) (E) for 5 days as described in Methods. The IFN-γ production from T-cell culture was detected by ELISA and statistically analyzed (n=3). (F) Statistically analyze of western blot of figure 5F (n=3). Results are representative of 3 independent experiments. *P<0.05, **P<0.01, and n.s. P>0.05 for groups connected by horizontal lines compared.

**S5 Fig. ADM-stimulated macrophages induce IFN-γ-producing T-cell responses.** (A) Secondary antibody-only stained sections as controls for the staining in Figure 6A. Scale bars: 50 microns. (B) IL-12p35 mRNA expression in gastric mucosa of uninfected donors (n=40) and *H. pylori*-infected patients (n=71) was compared. IL-12p35 mRNA expression in gastric mucosa of uninfected donors (n=40), *cagA*^-^ *H. pylori*-infected (n=24), and *cagA*+ *H. pylori*-infected (n=47) patients was compared. (C and D) The IFN-γ production from T cell-macrophage co-culture was detected by ELISA as described in Methods and statistically analyzed (n=3). *P<0.05, **P<0.01, and n.s., *P*>0.05 for groups connected by horizontal lines compared.

**S1 Table. Clinical characteristics of patients**

**S2 Table. Antibodies and other reagents**

**S3 Table. Primer and probe sequences for real-time PCR analysis**

## References

1. Hooi JKY, Lai WY, Ng WK, Suen mmy, Underwood FE, Tanyingoh D, et al. Global Prevalence of Helicobacter pylori Infection: Systematic Review and Meta-Analysis. Gastroenterology. 2017;153(2):420–429.

2. Suerbaum S, Michetti P. Helicobacter pylori infection. N Engl J Med. 2002;347(15):1175–1186.

3. Johnson KS, Ottemann KM. Colonization, localization, and inflammation: the roles of H. pylori chemotaxis in vivo. Curr Opin Microbiol. 2018;41:51–57.

4. Neumann L, Mueller M, Moos V, Heller F, Meyer TF, Loddenkemper C, et al. Mucosal Inducible NO Synthase-Producing IgA+ Plasma Cells in Helicobacter pylori-Infected Patients. J Immunol. 2016;197(5):1801–1808.

5. Wei L, Wang J, Liu Y. Prior to Foxp3(+) regulatory T-cell induction, interleukin-10-producing B cells expand after Helicobacter pylori infection. Pathog Dis. 2014;72(1):45–54.

6. Whitmore LC, Weems MN, Allen LH. Cutting Edge: Helicobacter pylori Induces Nuclear Hypersegmentation and Subtype Differentiation of Human Neutrophils In Vitro. J Immunol. 2017;198(5):1793–1797..

7. Hinson JP, Kapas S, Smith DM. Adrenomedullin, a multifunctional regulatory peptide. Endocr Rev. 2000;21(2):138–167.

8. Takei Y, Inoue K, Ogoshi M, Kawahara T, Bannai H, Miyano S, et al. Identification of novel adrenomedullin in mammals: a potent cardiovascular and renal regulator. FEBS Lett. 2004;556(1-3):53–58.

9. Fukuda K, Tsukada H, Oya M, Onomura M, Kodama M, Nakamura H, et al. Adrenomedullin promotes epithelial restitution of rat and human gastric mucosa in vitro. Peptides. 1999;20(1):127–132.

10. Groschl M, Wendler O, Topf HG, Bohlender J, Köhler H, et al. Significance of salivary adrenomedullin in the maintenance of oral health: stimulation of oral cell proliferation and antibacterial properties. Regul Pept. 2009;154(i1-3):16–22.

11. Hirsch AB, McCuen RW, Arimura A, Schubert ML. Adrenomedullin stimulates somatostatin and thus inhibits histamine and acid secretion in the fundus of the stomach. Regul Pept. 2003;110(3):189–195.

12. Martinez, A. A new family of angiogenic factors. Cancer Lett. 2006;236(2):157–163.

13. Nagata S, Yamasaki M, Kitamura K. Anti-Inflammatory Effects of PEGylated Human Adrenomedullin in a Mouse DSS-Induced Colitis Model. Drug Dev Res. 2017;78(3-4):129–134.

14. Bauer B, Pang E, Holland C, Kessler M, Bartfeld S, Meyer TF, et al. The Helicobacter pylori virulence effector CagA abrogates human beta-defensin 3 expression via inactivation of EGFR signaling. Cell Host Microbe. 2012;11(6):576–586.

15. Serrano C, Wright SW, Bimczok D, Shaffer CL, Cover TL, Venegas A, et al. Downregulated Th17 responses are associated with reduced gastritis in Helicobacter pylori-infected children. Mucosal Immunol. 2013;6(5):950–959.

16. Hitzler I, Oertli M, Becher B, Agger EM, Müller A. Dendritic cells prevent rather than promote immunity conferred by a helicobacter vaccine using a mycobacterial adjuvant. Gastroenterology. 2011;141(1):186–96, 196.e1.

17. Feng CG, D, Kullberg M, Cheever A, Scanga CA, Hieny S, et al. Maintenance of pulmonary Th1 effector function in chronic tuberculosis requires persistent IL-12 production. J Immunol. 2005;174(7):4185–4192.

18. Hickman SP, Chan J, Salgame P. Mycobacterium tuberculosis induces differential cytokine production from dendritic cells and macrophages with divergent effects on naive T cell polarization. J Immunol. 2002;168(9):4636–4642.

19. Beceiro S, Radin JN, Chatuvedi R, Piazuelo MB, Horvarth DJ, Cortado H, et al. TRPM2 ion channels regulate macrophage polarization and gastric inflammation during Helicobacter pylori infection. Mucosal Immunol. 2017 Mar;10(2):493–507.

20. Dorer MS, Talarico S, Salama NR. Helicobacter pylori’s unconventional role in health and disease. PLoS Pathog. 2009;5(10):e1000544.

21. Debowski AW, Walton SM, Chua EG, Tay AC, Liao T, Lamichhane B, et al. Helicobacter pylori gene silencing in vivo demonstrates urease is essential for chronic infection. PLoS Pathog. 2017;13(6):e1006464.

22. Gupta VR, Patel HK, Kostolansky SS, Ballivian RA, Eichberg J, Blanke SR. Sphingomyelin functions as a novel receptor for Helicobacter pylori VacA. PLoS Pathog. 2008;4(5):e1000073.

23. Stein SC, Faber E, Bats SH, Murillo T, Speidel Y, Coombs N, et al. Helicobacter pylori modulates host cell responses by CagT4SS-dependent translocation of an intermediate metabolite of LPS inner core heptose biosynthesis. PLoS Pathog. 2017;13(7):e1006514.

24. Kitamura K, Kangawa K, Kawamoto M, Ichiki Y, Nakamura S, Matsuo H, et al. Adrenomedullin: a novel hypotensive peptide isolated from human pheochromocytoma. Biochem Biophys Res Commun. 1993;192(2):553–560.

25. Martinez-Herrero S, Martinez A. Adrenomedullin regulates intestinal physiology and pathophysiology. Domest Anim Endocrinol. 2016;56 Suppl:S66–83.

26. Matheson PJ, Mays MP, Hurt RT, Harris PD, Garrison RN. Adrenomedullin is increased in the portal circulation during chronic sepsis in rats. Am J Surg. 2003;186(5):519–525.

27. Allaker RP, Kapas S. Adrenomedullin and mucosal defence: interaction between host and microorganism. Regul Pept. 2003;112(1-3):147–152.

28. Allaker RP, Kapas S. Adrenomedullin expression by gastric epithelial cells in response to infection. Clin Diagn Lab Immunol. 2003;10(4):546–551.

29. Nishimatsu H, Suzuki E, Nagata D, Moriyama N, Satonaka H, Walsh K, et al. Adrenomedullin induces endothelium-dependent vasorelaxation via the phosphatidylinositol 3-kinase/Akt-dependent pathway in rat aorta. Circ Res. 200;89(1):63–70.

30. Fritz-Six kl, Dunworth WP, Li M, Caron KM. Adrenomedullin signaling is necessary for murine lymphatic vascular development. J Clin Invest. 2008;118(1):40–50.

31. Ichikawa-Shindo Y, Sakurai T, Kamiyoshi A, Kawate H, Iinuma N, Yoshizawa T, et al. The GPCR modulator protein RAMP2 is essential for angiogenesis and vascular integrity. J Clin Invest. 2008;118(1):29–39.

32. Koyama T, Ochoa-Callejero L, Sakurai T, Kamiyoshi A, Ichikawa-Shindo Y, Iinuma N, et al. Vascular endothelial adrenomedullin-RAMP2 system is essential for vascular integrity and organ homeostasis. Circulation. 2013;127(7):842–853.

33. Dackor R, Fritz-Six K, Smithies O, Caron K. Receptor activity-modifying proteins 2 and 3 have distinct physiological functions from embryogenesis to old age. J Biol Chem. 2007;282(25):18094–18099.

34. Liverani E, McLeod JD, Paul C. Adrenomedullin receptors on human T cells are glucocorticoid-sensitive. Int Immunopharmacol. 2012;14(1):75–81.

35. Rulle S, Ah Kioon md, Asensio C, Mussard J, Ea HK, Boissier MC, et al. Adrenomedullin, a neuropeptide with immunoregulatory properties induces semi-mature tolerogenic dendritic cells. Immunology. 2012;136(2):252–264.

36. Takano S, Uchida K, Miyagi M, Inoue G, Aikawa J, Iwabuchi K, et al. Adrenomedullin Regulates IL-1beta Gene Expression in F4/80+ Macrophages during Synovial Inflammation. J Immunol Res. 2017;2017:9832430.

37. Ismail HF, Fick P, Zhang J, Lynch RG, Berg DJ, et al. Depletion of neutrophils in IL-10(-/-) mice delays clearance of gastric Helicobacter infection and decreases the Th1 immune response to Helicobacter. J Immunol. 2003;170(7):3782–3789.

38. Krueger S, Kuester D, Bernhardt A, Wex T, Roessner A. Regulation of cathepsin X overexpression in H. pylori-infected gastric epithelial cells and macrophages. J Pathol. 2009;217(4):581–588.

39. Otani K, Tanigawa T, Watanabe T, Nadatani Y, Sogawa M, Yamagami H, et al. Toll-like receptor 9 signaling has anti-inflammatory effects on the early phase of Helicobacter pylori-induced gastritis. Biochem Biophys Res Commun. 2012;426(3):342–349.

40. Garhart CA, Redline RW, Nedrud JG, Czinn SJ. Clearance of Helicobacter pylori Infection and Resolution of Postimmunization Gastritis in a Kinetic Study of Prophylactically Immunized Mice. Infect Immun. 2002;70(7):3529–3538.

41. Roussel Y, Wilks M, Harris A, Mein C, Tabaqchali S. Evaluation of DNA extraction methods from mouse stomachs for the quantification of H. pylori by real-time PCR. J Microbiol Methods. 2005;62(1):71–81..

42. Mikula M, Dzwonek A, Jagusztyn-Krynicka K, Ostrowski J. Quantitative detection for low levels of Helicobacter pylori infection in experimentally infected mice by real-time PCR. J Microbiol Methods. 2003;55(2):351–359.

43. Wang TT, Zhao YL, Peng LS, Chen N, Chen W, Lv YP, et al. Tumour-activated neutrophils in gastric cancer foster immune suppression and disease progression through GM-CSF-PD-L1 pathway. Gut. 2017;66(11):1900–1911.

